# Human cytomegalovirus mediates APOBEC3B relocalization early during infection through a ribonucleotide reductase-independent mechanism

**DOI:** 10.1101/2023.01.30.526383

**Authors:** Elisa Fanunza, Adam Z. Cheng, Ashley A. Auerbach, Bojana Stefanovska, Sofia N. Moraes, James R. Lokensgard, Matteo Biolatti, Valentina Dell’Oste, Craig J. Bierle, Wade A. Bresnahan, Reuben S. Harris

## Abstract

The APOBEC3 family of DNA cytosine deaminases comprises an important arm of the innate antiviral defense system. The gamma-herpesviruses EBV and KSHV and the alpha-herpesviruses HSV-1 and HSV-2 have evolved an efficient mechanism to avoid APOBEC3 restriction by directly binding to APOBEC3B and facilitating its exclusion from the nuclear compartment. The only viral protein required for APOBEC3B relocalization is the large subunit of the ribonucleotide reductase (RNR). Here, we ask whether this APOBEC3B relocalization mechanism is conserved with the beta-herpesvirus human cytomegalovirus (HCMV). Although HCMV infection causes APOBEC3B relocalization from the nucleus to the cytoplasm in multiple cell types, the viral RNR (UL45) is not required. APOBEC3B relocalization occurs rapidly following infection suggesting involvement of an immediate early or early (IE-E) viral protein. In support of this mechanism, cycloheximide treatment of HCMV-infected cells prevents the expression of viral proteins and simultaneously blocks APOBEC3B relocalization. In comparison, the treatment of infected cells with phosphonoacetic acid, which is a viral DNA synthesis inhibitor affecting late protein expression, still permits A3B relocalization. These results combine to show that the beta-herpesvirus HCMV uses a fundamentally different, RNR-independent molecular mechanism to antagonize APOBEC3B.

**Importance:** Human cytomegalovirus (HCMV) infections can range from asymptomatic to severe, particularly in neonates and immunocompromised patients. HCMV has evolved strategies to overcome host-encoded antiviral defenses in order to achieve lytic viral DNA replication and dissemination and, under some conditions, latency and long-term persistence. Here, we show that HCMV infection causes the antiviral factor, APOBEC3B, to relocalize from the nuclear compartment to the cytoplasm. This overall strategy resembles that used by related herpesviruses. However, the HCMV relocalization mechanism utilizes a different viral factor(s) and available evidence suggests the involvement of at least one protein expressed at the early stages of infection. This knowledge is important because a greater understanding of this mechanism could lead to novel antiviral strategies that enable APOBEC3B to naturally restrict HCMV infection.

## Introduction

The APOBEC3 (A3) system is an essential part of the cellular innate immune response to viral infections [reviewed by (1–3)]. A3-mediated restriction has been reported for a broad number of DNA-based viruses, including exogenous viruses (retroviruses, polyomaviruses, papillomaviruses, parvoviruses, hepadnaviruses, and herpesviruses) and endogenous viruses and transposable elements. The mechanism by which virus restriction occurs is well-documented and dependent partly on the ability of A3 enzymes to introduce mutations in the viral genome by catalyzing cytosine deamination in exposed single stranded (ss)DNA intermediates. In addition, deaminase-independent antiviral activity has been reported against endogenous retroelements, reverse-transcribing viruses, adeno-associated viruses, and RNA viruses, and this may be attributed to strong nucleic acid binding activity.

The continuous arms race between host and viruses leads to the selection of viral factors able to counteract innate immune factors, including the A3 antiviral enzymes. For example, HIV-1, HIV-2, and related lentiviruses encode a viral accessory protein Vif that mediates the degradation of restrictive A3s (4, 5). Recently, a novel mechanism of A3 counteraction was discovered for the gamma-herpesviruses Epstain-Barr virus (EBV), which use the viral ribonucleotide reductase (RNR) large subunit, BORF2, to directly bind, inhibit, and relocalize APOBEC3B (A3B) from the nucleus to the cytoplasm, thus preserving viral genome integrity (6). This mechanism of A3 neutralization is likely to be conserved because at least two other herpesviruses, Kaposi’s sarcoma-associated herpesvirus (KSHV), and herpes simplex virus 1 (HSV-1), whose RNRs (ORF61, and ICP6, respectively) physically interact with A3B, as well with APOBEC3A (A3A), and trigger their redistribution from the nucleus to the cytoplasmic compartment (7–10). In further support of evolutionary conservation, a systematic analysis of a large panel of present-day gamma-herpesvirus RNRs and primate A3B proteins indicates that the evolution of this viral RNR-mediated A3B neutralization mechanism was likely selected by the birth of the *A3B* gene by unequal crossing-over in an ancestral Old World primate approximately 29-43 million years ago (8, 11).

Human cytomegalovirus (HCMV) is a member of the beta-herpesvirus subfamily. HCMV is a ubiquitous virus, found in approximately 90% of the worldwide population. HCMV infection is usually asymptomatic in healthy individuals, but it can cause severe disease in immunocompromised hosts [reviewed by (12, 13)]. Congenital HCMV infections are also a leading cause of birth defects [reviewed by (14, 15)]. HCMV has a large double-stranded (ds)DNA genome of 235 kb – the largest among known human herpesviruses – containing 165 canonical open reading frames (ORFs) and several alternative transcripts [reviewed by (16)]. Lytic HCMV infection involves a temporal cascade of gene expression. A small subset of genes, termed immediate-early genes (IE), are the first to be expressed. Transcription of IE genes does not require *de novo* protein synthesis. Immediate-early proteins together with host factors mediate the expression of the kinetically distinct early genes (E), whose products in large part promote viral genome replication and the expression of late genes (L) [reviewed by (16)].

Several HCMV gene products have acquired the ability to subvert different signaling pathways and modulate various components of the immune response to make the host cellular machinery more permissible to viral replication and survival [reviewed by (17, 18)]. Given the ability of gamma- and alpha-herpesviruses (EBV/KSHV and HSV-1/2, respectively) to inhibit A3B, we sought to investigate whether HCMV possesses a similar RNR-mediated A3 neutralization mechanism. Our results demonstrate that HCMV infection is also capable of inducing the selective nuclear to cytoplasmic relocalization of A3B. However, surprisingly, results with multiple independent viral strains and cell lines indicate that the relocalization mechanism of A3B by HCMV is not conserved with other human herpesviruses and, instead, occurs independently of the HCMV UL45 RNR. In addition to this strong mechanistic distinction, multiple lines of evidence including rapid A3B relocalization kinetics suggest involvement of at least one viral IE-E protein in A3B relocalization.

## Results

### HCMV mediates A3B relocalization independently of viral strain and cell type

We previously reported the ability of gamma- and alpha-herpesviruses to bind to A3B and mediate its relocalization from the nuclear compartment into cytoplasmic aggregates (6–8). To investigate whether the beta-herpesvirus HCMV has similar functionality, immunofluorescence (IF) microscopy experiments were done using infected primary human foreskin fibroblast-1 (HFF-1) cells. First, HFF-1 cells were stably transduced with a lentivirus expressing C-terminally HA-tagged A3B. As reported for other human cell types (11, 19–21), A3B localizes primarily to the nuclear compartment of mock/non-infected HFF-1 cells (representative image in **Fig. 1A** and quantification in **Fig. 1F**). Next, HFF-1 transduced cells were infected with HCMV strain TB40/E that expresses the mCherry protein (TB40-mCherry) and analyzed for A3B localization by IF microscopy 72 hpi. Infected, mCherry-positive cells are visibly enlarged, as expected for productive cytomegalovirus infection, and A3B becomes predominantly cytoplasmic (representative image in **Fig. 1A** and quantification in **Fig. 1F**). When HFF-1 cells were transduced to express the other seven human A3 family members, A3B is the only protein to show a major change in subcellular distribution (**Supplementary Fig. 1**).

**Fig 1.**
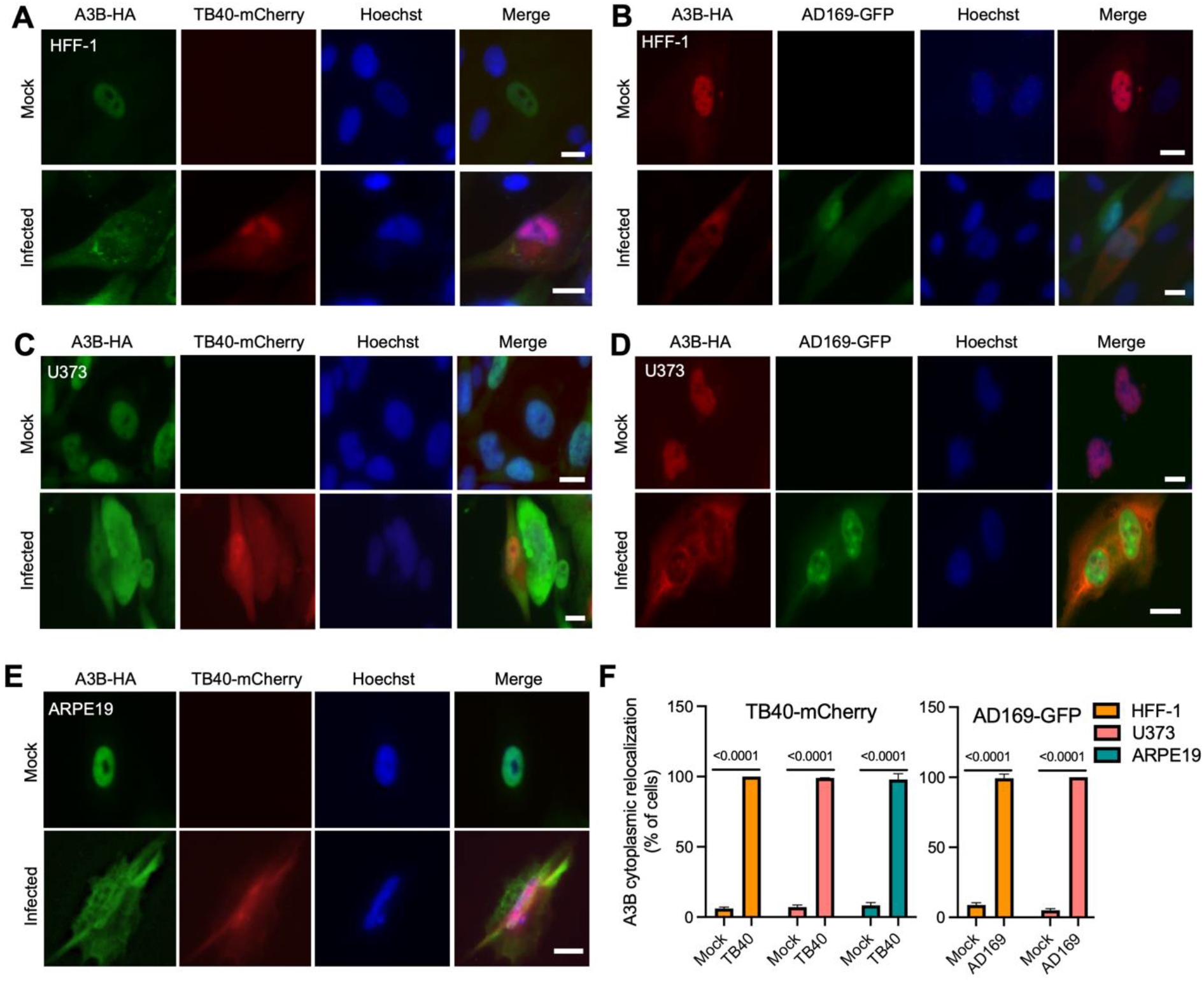
A3B relocalization occurs with multiple HCMV strains in different cell types. (**A-E**) Representative IF microscopy images of the indicated cell types stably expressing A3B-HA incubated with medium alone (mock) or infected with the indicated HCVM strains for 72 hrs (10 μm scale). (**F**) Quantification A3B-HA subcellular localization phenotypes shown in panels A-E. Each histogram bar reports the percentage of cells with cytoplasmic A3B-HA (n>100 cells per condition; mean +/− SD with indicated p-values from unpaired student’s t-tests).

To ask if the A3B relocalization mechanism extends to other HCMV strains, HFF-1 stably transduced with HA-tagged A3B were infected with the laboratory-adapted GFP-expressing AD169 strain (AD169-GFP), and IF microscopy was done 72 hpi. As above with TB40-mCherry, AD169-GFP infection induces strong relocalization of A3B from the nuclear compartment to the cytoplasm (representative image in **Fig. 1B** and quantification in **Fig. 1F**). Similar A3B-HA relocalization is observed during infection of HFF-1 cells with the Merlin strain (**Supplementary Fig. 2A**). A3B-HA relocalization is also observed in other cell types including the human glioma cell line, U373, infected with TB40-mCherry (**Fig. 1C**) or AD169-GFP (**Fig. 1D**) and the human retinal pigment epithelial cells, ARPE19, infected with TB40-mCherry (**Fig. 1E,** and **1F**). As an additional control for specificity, A3B-EGFP but not EGFP alone relocalizes to the cytoplasmic compartment following infection of ARPE19 cells with TB40-mCherry (**Supplementary Fig. 2B**). Thus, the A3B relocalization phenotype is evident following infection with multiple HCMV strains and in a range of different cell types (both primary and immortalized) permissive for HCMV infection.

### Catalytic mutant and endogenous A3B are relocalized upon HCMV infection

Overexpression of wildtype A3B causes chromosomal DNA deamination, strong DNA damage responses, cell cycle perturbations, and eventually cell death (22–24). These phenotypes require the catalytic activity of A3B. To address the possibility that A3B relocalization may be triggered indirectly by one of these events, HFF-1, U373, and ARPE19 cells were transduced with a lentiviral construct expressing the catalytically inactive A3B mutant (A3B-E255A). HCMV infection and IF microscopy experiments were done as above. In all instances, A3B-E255A relocalizes from the nucleus to the cytoplasm following infection with TB40-mCherry or AD169-GFP (representative images in **Fig. 2A-E** and quantification in **Fig. 2F**). These results demonstrate that the relocalization of A3B occurs independent of its DNA deamination activity and is unlikely to be part of a general DNA damage response.

**Fig. 2.**
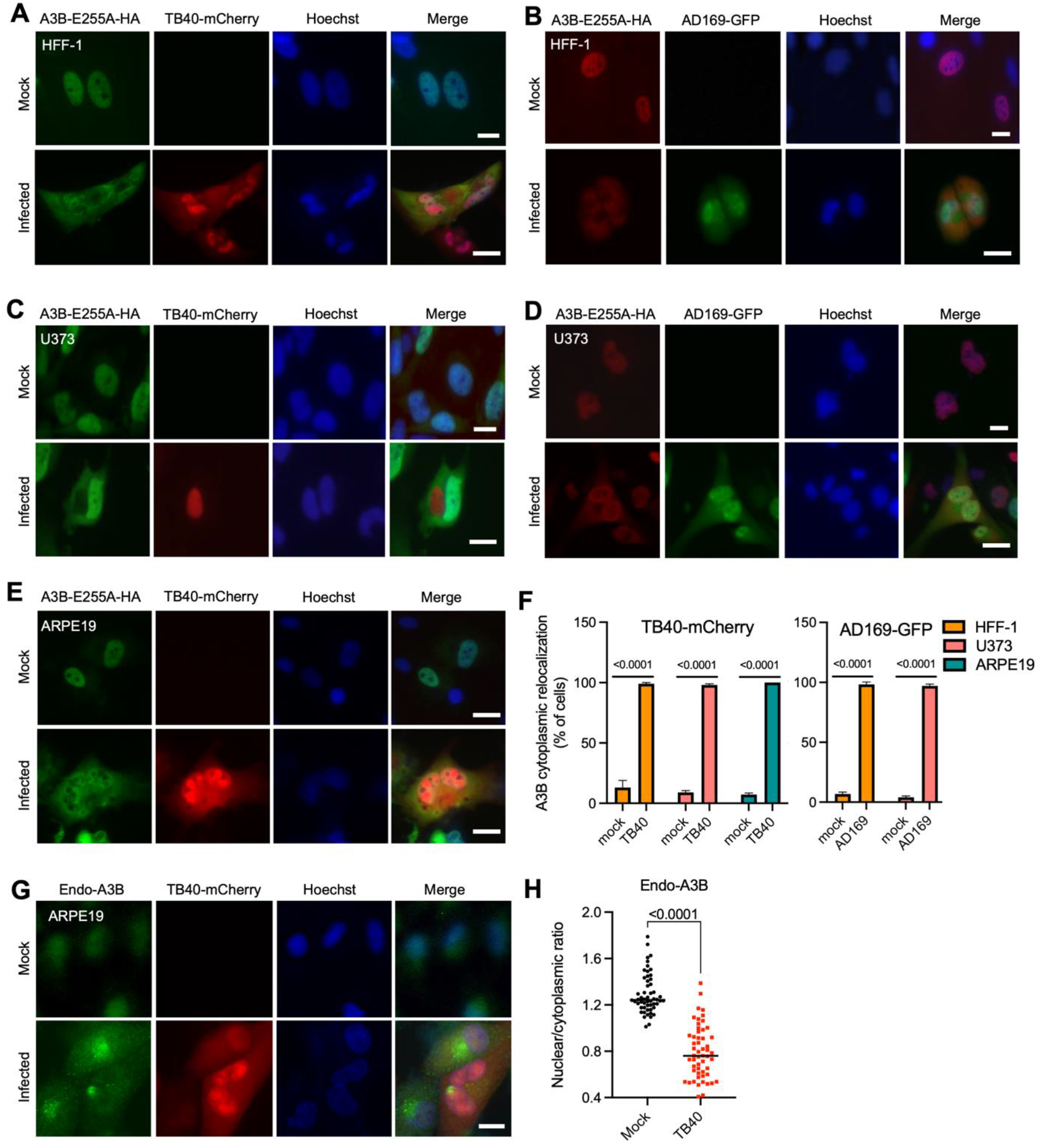
Catalytic mutant and endogenous A3B are relocalized by HCMV. (**A-E**) Representative IF microscopy images of the indicated cell types stably expressing A3B-E255A-HA incubated with medium alone (mock) or infected with the indicated HCVM strains for 72 hrs (10 μm scale). (**F**) Quantification A3B-E255A-HA subcellular localization phenotypes shown in panels A-E. Each histogram bar reports the percentage of cells with cytoplasmic A3B-HA (n>100 cells per condition; mean +/− SD with indicated p-values from unpaired student’s t-tests). (**G**) Representative IF microscopy images of ARPE19 cells incubated with medium alone (mock) or infected with TB40-mCherry for 72 hrs, stained for endogenous A3B (10 μm scale). (**H**) Quantification of endogenous A3B subcellular localization phenotype shown in panel G. The dot-plot chart shows the ratio between nuclear and cytoplasmic fluorescence intensity (n>50 cells per condition; p-values were obtained using unpaired student’s t-tests).

To further confirm that the relocalization phenotype is not a general effect of A3B overexpression, we next evaluated the subcellular localization of the endogenous protein. ARPE19 cells were infected with TB40-mCherry, allowing 72 hrs for infection to progress, and then performing IF microscopy with the rabbit anti-human A3B monoclonal antibody 5210-87-13 (25). As observed above with overexpressed A3B-HA with or without catalytic activity, the endogenous A3B protein also shows strong relocalization from the nucleus to the cytoplasm (**Fig. 2G-H**). These results combine to indicate that the A3B relocalization mechanism of HCMV is deamination-independent and not likely to be an artifact of protein overexpression because endogenous A3B also has a clear phenotype.

### HCMV UL45 is incapable of binding, inhibiting, or relocalizing human A3B

The only gamma- and alpha-herpesvirus protein required for A3B relocalization is the large subunit of the viral RNR (6–8). The large RNR subunit of EBV, BORF2, directly binds A3B, inhibits its catalytic activity, and relocalizes the protein from the nucleus to the cytoplasm. To address whether the HCMV large RNR subunit, UL45, is capable of similarly binding to A3B, we performed a series of coimmunoprecipitation (co-IP) experiments. HEK293T cells were transfected with empty vector or FLAG-tagged HCMV UL45 or EBV BORF2 together with a HA-tagged human A3B or other A3 constructs as negative controls. As expected, EBV BORF2 robustly co-IPs A3B but not A3G (**Fig. 3A**). In parallel experiments, HCMV UL45 appears incapable of co-IP of either A3B or A3A (**Fig. 3A**). However, conclusions from these experiments are limited by relatively low UL45 expression levels in cell extracts, multiple expressed products including likely monomeric and dimeric forms (full-length UL45 is predicted to be ~ 108 kDa), and lack of a positive control for UL45 co-IP (HCMV lacks a small RNR subunit that normally associates with the large RNR subunit and UL45 interactors have yet to be reported).

**Fig. 3.**
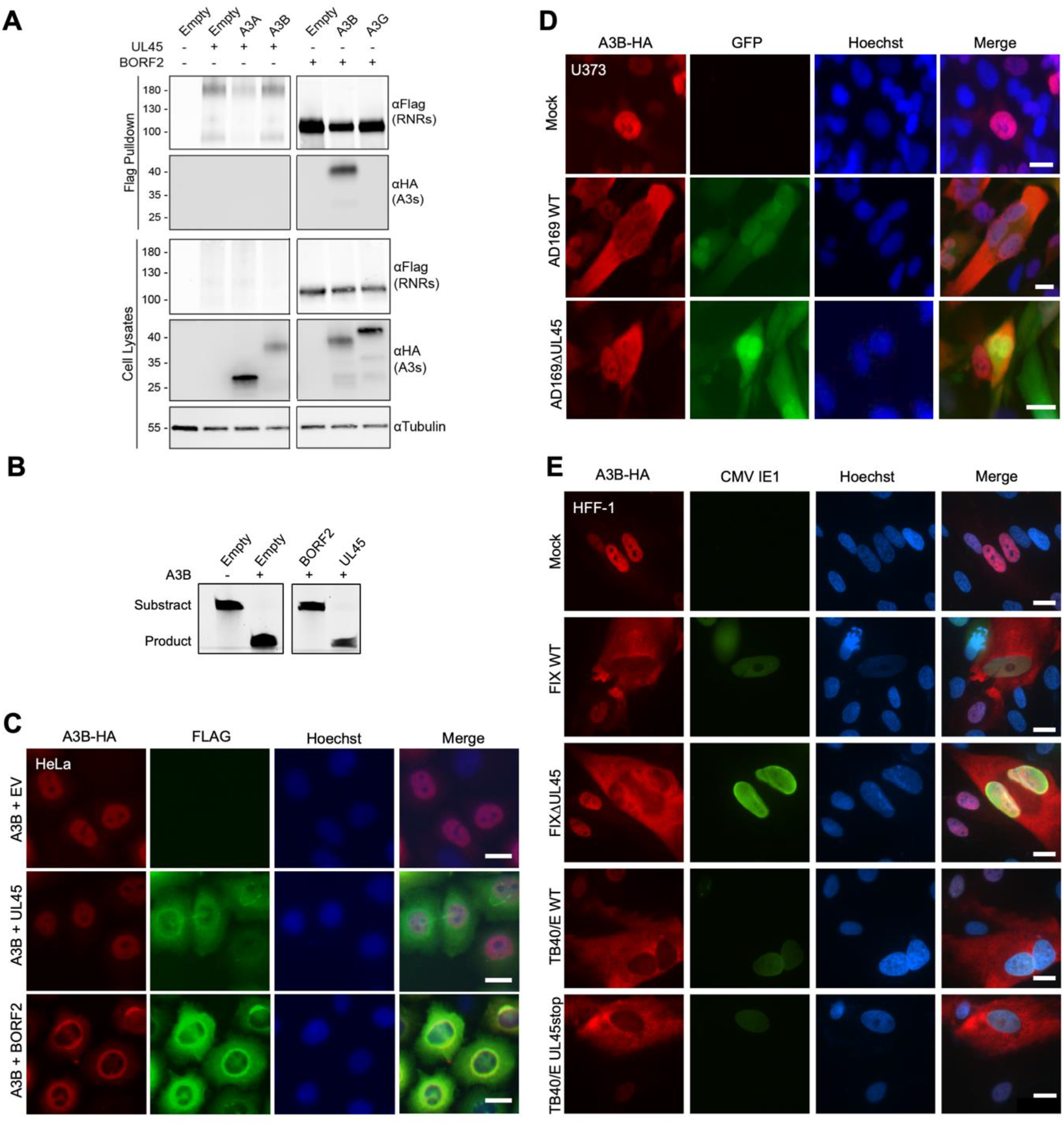
A3B relocalization is UL45-independent. (**A**) Coimmunoprecipitation of transfected HCMV UL45-FLAG with the indicated A3-HA constructs in HEK293T cells. Cells co-transfected with EBV BORF2 and A3B or A3G are used as positive and negative controls, respectively. (**B**) TBE-urea PAGE analysis of A3B deaminase activity in the presence of empty vector, HCMV UL45, or EBV BORF2. (**C**) Representative IF microscopy images of HeLa cells transiently expressing A3B-HA together with empty vector, HCMV UL45-FLAG or EBV BORF2-FLAG (10 μm scale). (**D-E**) Representative IF microscopy images of the indicated cell types stably expressing A3B-HA incubated with medium alone (mock) or infected with the indicated HCVM strains and UL45-null derivatives for 72 hrs (10 μm scale).

We, therefore, turned to other approaches to ask whether HCMV UL45 is capable of interfering with A3B catalytic activity. First, single-stranded (ss)DNA C-to-U activity assays were completed using extracts from HEK293T cells expressing viral RNR large subunits and A3B. Consistent with previous results (6), A3B exhibits robust ssDNA C-to-U activity in cell extracts and its activity is strongly inhibited by BORF2 (**Fig. 3B**). In comparison, HCMV UL45 co-expression has a negligible effect on the ssDNA C-to-U activity of A3B in cell extracts (**Fig. 3B**). Next, IF microscopy experiments were done by cotransfecting HeLa cells with A3B-HA and viral RNR-FLAG constructs, allowing 48 hrs for expression, and imaging with specific antibodies. In contrast to EBV BORF2, which relocalizes A3B from the nuclear to the cytoplasmic compartment, expression of HCMV UL45 has no effect on A3B subcellular localization (**Fig. 3C**). Taken together, negative results from co-IP, deaminase inhibition, and colocalization experiments indicate that the large RNR subunit of HCMV, UL45, is incapable of interacting with A3B and/or promoting its relocalization.

To directly ask whether HCMV UL45 is required for A3B relocalization, we compared the subcellular localization phenotypes of A3B in U373 cells following infection by AD169-GFP or a derivative virus engineered to lack UL45 [AD169-GFP ΔUL45 (26)]. U373 cells were stably transduced with HA-tagged A3B 48 hrs prior to mock infection or infection with AD169-GFP or AD169-GFP ΔUL45. After 72 hrs of infection, cells were fixed, permeabilized, and imaged by IF microscopy. As described above, infection by AD169-GFP causes the relocalization of A3B from the nuclear to the cytoplasmic compartment (**Fig. 3D**). As expected, cells infected with AD169-GFP ΔUL45 show an indistinguishable A3B cytoplasmic relocalization phenotype (**Fig. 3D**). This key result was confirmed by IF microscopy experiments using two other HCMV strains (TB40/E and FIX) and otherwise isogenic UL45-null derivatives (TB40/E ΔUL45 and FIX ΔUL45) (**Fig. 3E**). These results demonstrate that UL45 is dispensable for HCMV-mediated relocalization of A3B and, together with the results above, that this beta-herpesvirus does not share the RNR-dependent mechanism of A3B relocation of the gamma- and alpha-herpesviruses.

### The N-terminal domain of A3B is sufficient for HCMV-mediated relocalization

A3B is comprised of two conserved cytidine deaminase domains: an inactive N-terminal domain (A3B-NTD) and a catalytically active C-terminal domain (A3B-CTD) (9, 27). A3B-NTD is thought to be regulatory in nature and is alone sufficient for nuclear localization (11, 19). EBV BORF2 mediates A3B relocalization by binding to the CTD and not the NTD (6). To ask whether domain requirements might further distinguish the A3B relocalization mechanism of HCMV, IF microscopy experiments were done with ARPE19 cells transfected with EGFP-tagged full-length A3B (A3B-FL), A3B-NTD, or A3B-CTD constructs. After 72 hrs infection with TB40-mCherry, A3B-FL shows clear relocalization to the cytoplasmic compartment in comparison to the unchanged cell-wide EGFP control (**Fig. 4A-B**). Surprisingly, A3B-NTD, which shows nuclear localization in mock-infected cells, becomes predominantly cytoplasmic after infection (**Fig. 4A-B**). A3B-CTD has a cell-wide localization pattern that is not changed by virus infection (**Fig. 4A-B**). In contrast, EBV BORF2 has no effect on A3B-NTD nuclear localization, and it strongly promotes the relocalization of A3B-FL and A3B-CTD into cytoplasmic aggregates (**Fig. 4C**). These data combine to show that A3B-NTD is sufficient for A3B subcellular redistribution during HCMV infection and additionally distinguish the molecular mechanism from that mediated by the large RNR protein of gamma- and alpha-herpesvirus.

**Fig. 4.**
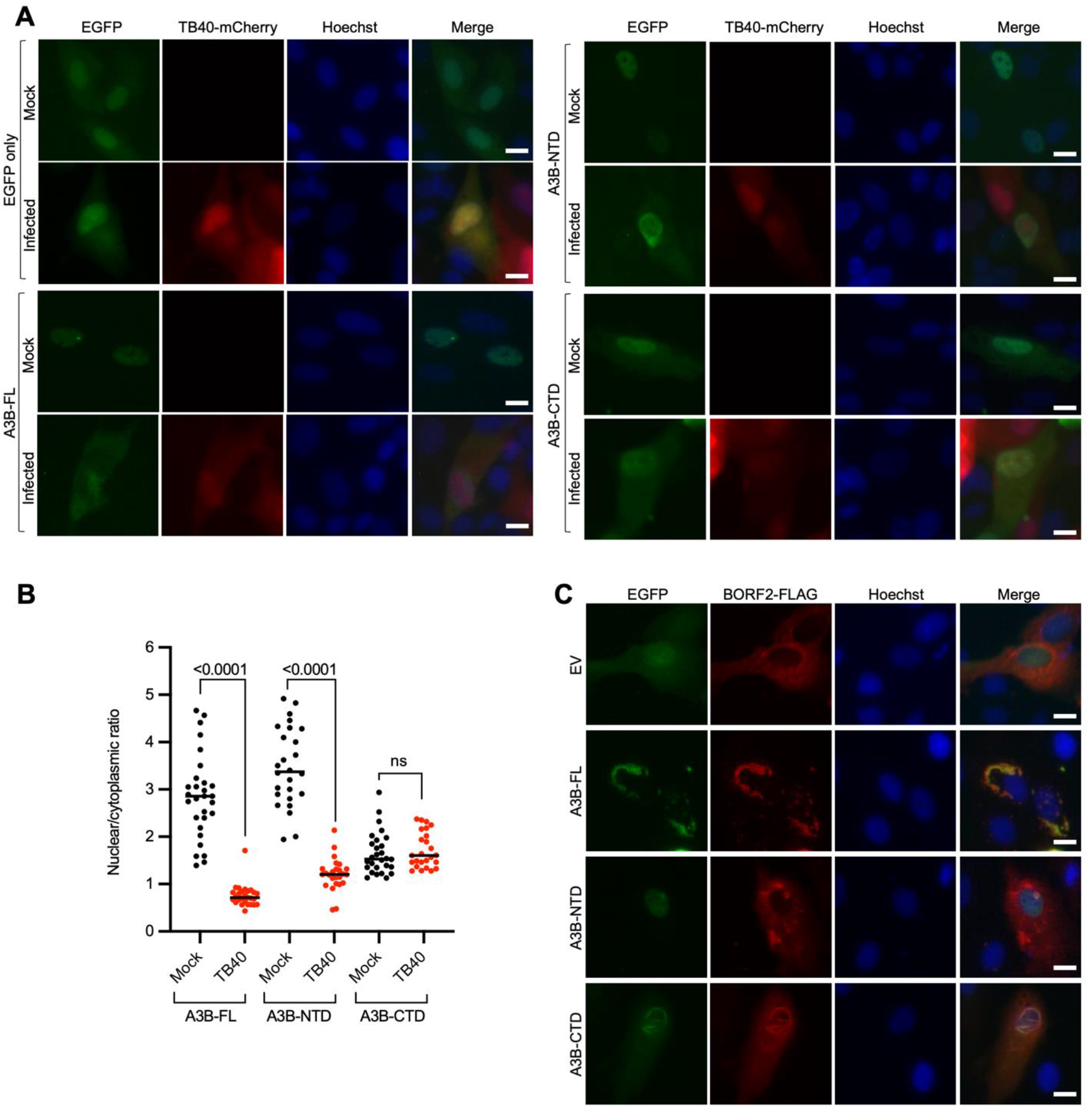
The NTD of A3B is sufficient for A3B relocalization mediated by HCMV. (**A**) Representative IF microscopy images of ARPE19 cells transiently expressing EGFP alone, A3B-FL-EGFP, A3B-NTD-EGFP, and A3B-CTD-EGFP, incubated with medium alone (mock) or infected with TB40-mCherry for 72 hrs (10 μm scale). (**B**) Quantification of A3B-FL, A3B-NTD, and A3B-CTD subcellular localization phenotype shown in panel A. The dot-plot chart shows the ratio between nuclear and cytoplasmic fluorescence intensity (n>25 cells per condition; p-values were obtained using unpaired student’s t-tests). (**C**) Representative IF microscopy images of HeLa cells transiently expressing EBV BORF2-FLAG together with EGFP alone, A3B-FL-EGFP, A3B-NTD-EGFP, and A3B-CTD-EGFP (10 μm scale).

### A3B relocalization occurs early during infection and requires *de novo* HCMV protein expression

During a productive HCMV infection, viral genes are expressed chronologically in three main groups (28). Immediate-early (IE) genes are are first expressed at between 2 and 6 hpi, early (E) genes are turned on between 4 and 12 hpi, and late (L) genes begin to express after after ~24 hpi and following the onset of viral DNA replication. To investigate the kinetics of A3B relocalization during HCMV infection, HFF-1 cells stably expressing A3B-HA were infected with TB40-mCherry or AD169-GFP and IF microscopy was performed at multiple timepoints after infection (6, 24, 48, and 72 hpi; **Fig. 5A-B** and **5C-D**, respectively). This experiment shows that relocalization begins to occur rapidly with most infected cells exhibiting partial or full A3B-HA relocalization at the earliest 6 hpi timepoint. Moreover, the percentage of cells exhibiting cytoplasmic A3B-HA increases over time and is complete by 72 hpi. These kinetics suggest that a HCMV IE or E protein may be responsible for A3B relocalization during infection.

**Fig. 5.**
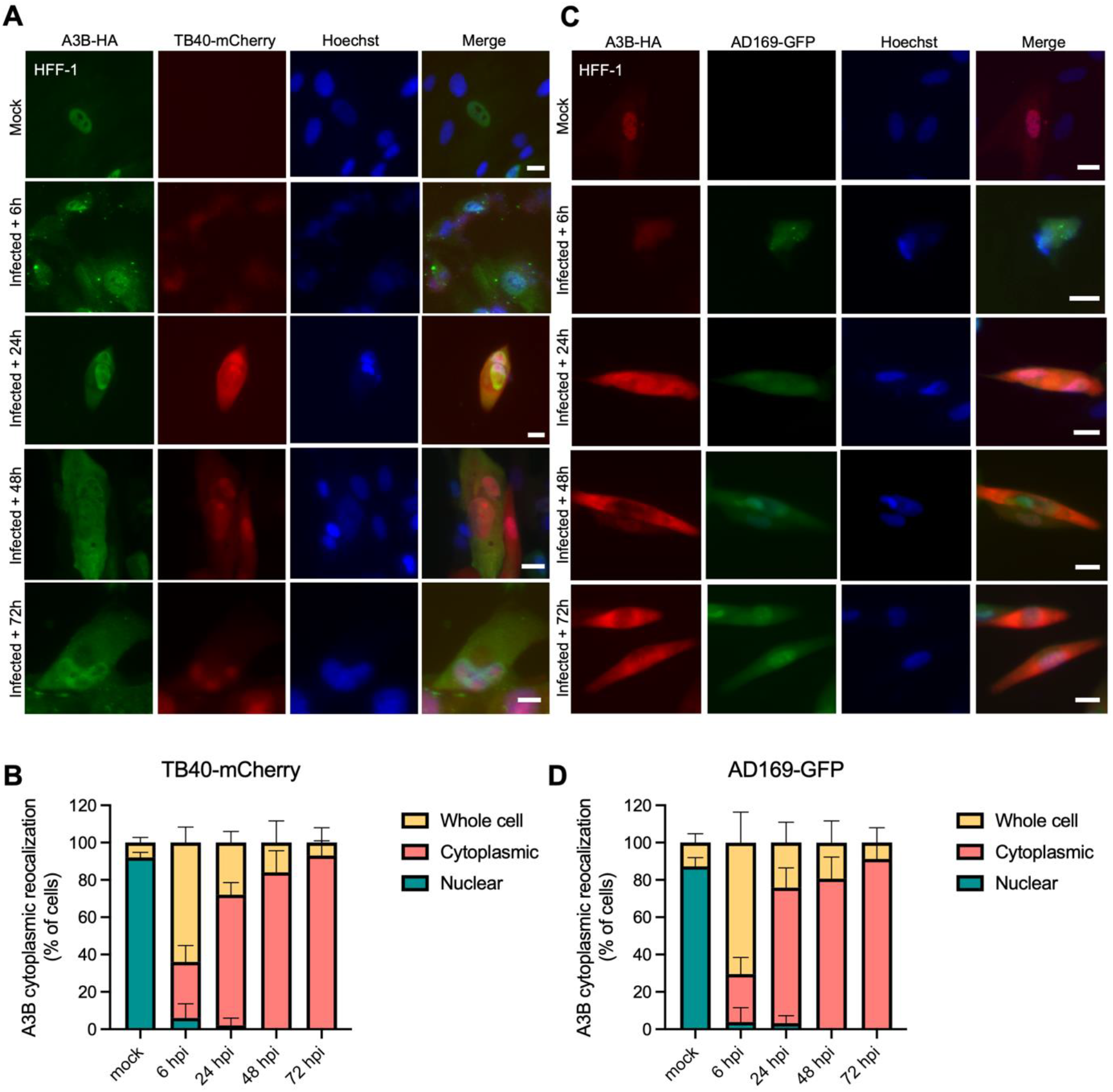
A3B relocalization occurs early during HCMV infection. (**A,C**) Representative IF microscopy images of HFF-1 cells stably expressing A3B-HA incubated with medium alone (mock) or infected with the indicated HCVM strains for the indicated time points (10 μm scale). (**B, D**) Quantification A3B-HA subcellular localization phenotypes shown in panels A and C. Each histogram bar reports the percentage of cells with whole cell, cytoplasmic, and nuclear A3B-HA (n>100 cells per condition; mean +/− SD with indicated p-values from unpaired student’s t-tests).

To further investigate whether *de novo* viral protein expression is required for A3B relocalization, HFF-1 cells stably expressing A3B-HA were infected with AD169-GFP, treated for 24 hrs with the translation inhibitor cycloheximide (CHX) or DMSO as a control, and then subjected to IF microscopy (**Fig. 6A**). CHX treatment strongly prevents A3B-HA from relocalizing to the cytoplasm, whereas DMSO treatment does not (**Fig. 6B** and **6E**). Similarly, cells infected with a recombinant AD169 lacking expression of the IE1 protein (AD169ΔIE1), is completely defective in A3B relocalization (**Fig. 6C**). In contrast, treating infected cells with phosphonoacetic acid (PAA), which blocks viral DNA synthesis and therefore also L protein expression, also fails to block A3B relocalization (**Fig. 6D** and **6E**). Taken together with the rapid relocalization kinetics described above, these additional experiments strongly implicate at least one HCMV IE/E protein in the A3B relocalization mechanism (**Fig. 7**).

**Fig. 6.**
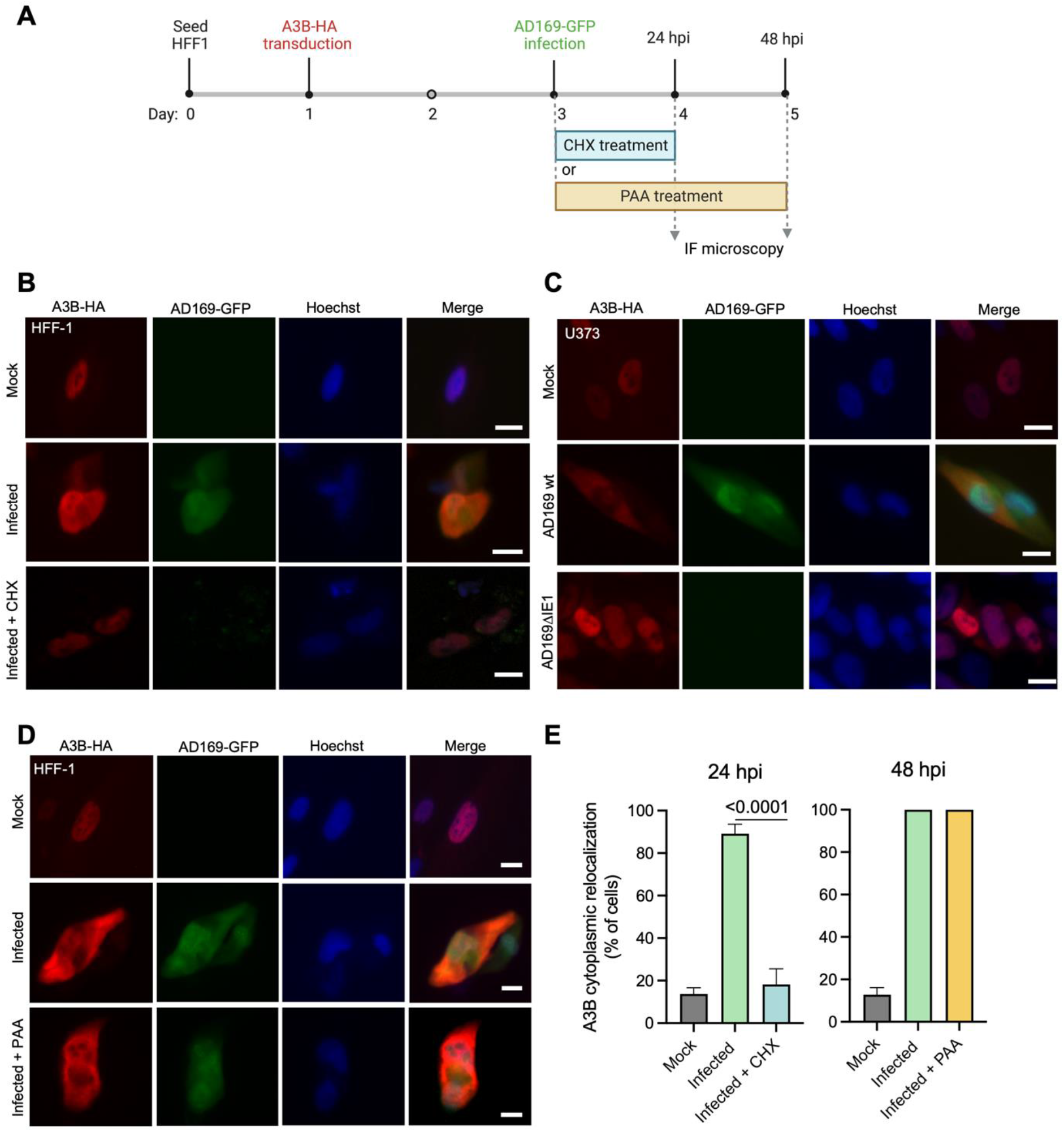
A3B relocalization requires *de novo* translation of HCMV proteins but it does not require viral DNA synthesis. (**A**) Schematic representation of cycloheximide (CHX) and phosphonoacetic acid (PAA) treatment in infected cells. Image created with BioRender. (**B**) Representative IF microscopy images of HFF-1 cells stably expressing A3B-HA incubated with medium alone (mock) infected with AD169-GFP and treated with DMSO or CHX for 24 hrs (10 μm scale). (**C**) Representative IF microscopy images of HFF-1 cells stably expressing A3B-HA incubated with medium alone (mock) infected with AD169-GFP, or AD169-GFP ΔIE1 for 72 hrs (10 μm scale). (**D**) Representative IF microscopy images of HFF-1 cells stably expressing A3B-HA incubated with medium alone (mock) or infected with AD169-GFP and treated with DMSO or PAA for 48 hrs (10 μm scale). (**E**) Quantification A3B-HA subcellular localization phenotypes shown in panels B and D. Each histogram bar reports the percentage of cells with cytoplasmic A3B-HA (n>80 cells per condition; mean +/− SD with indicated p-values from unpaired student’s t-tests).

**Fig 7.**
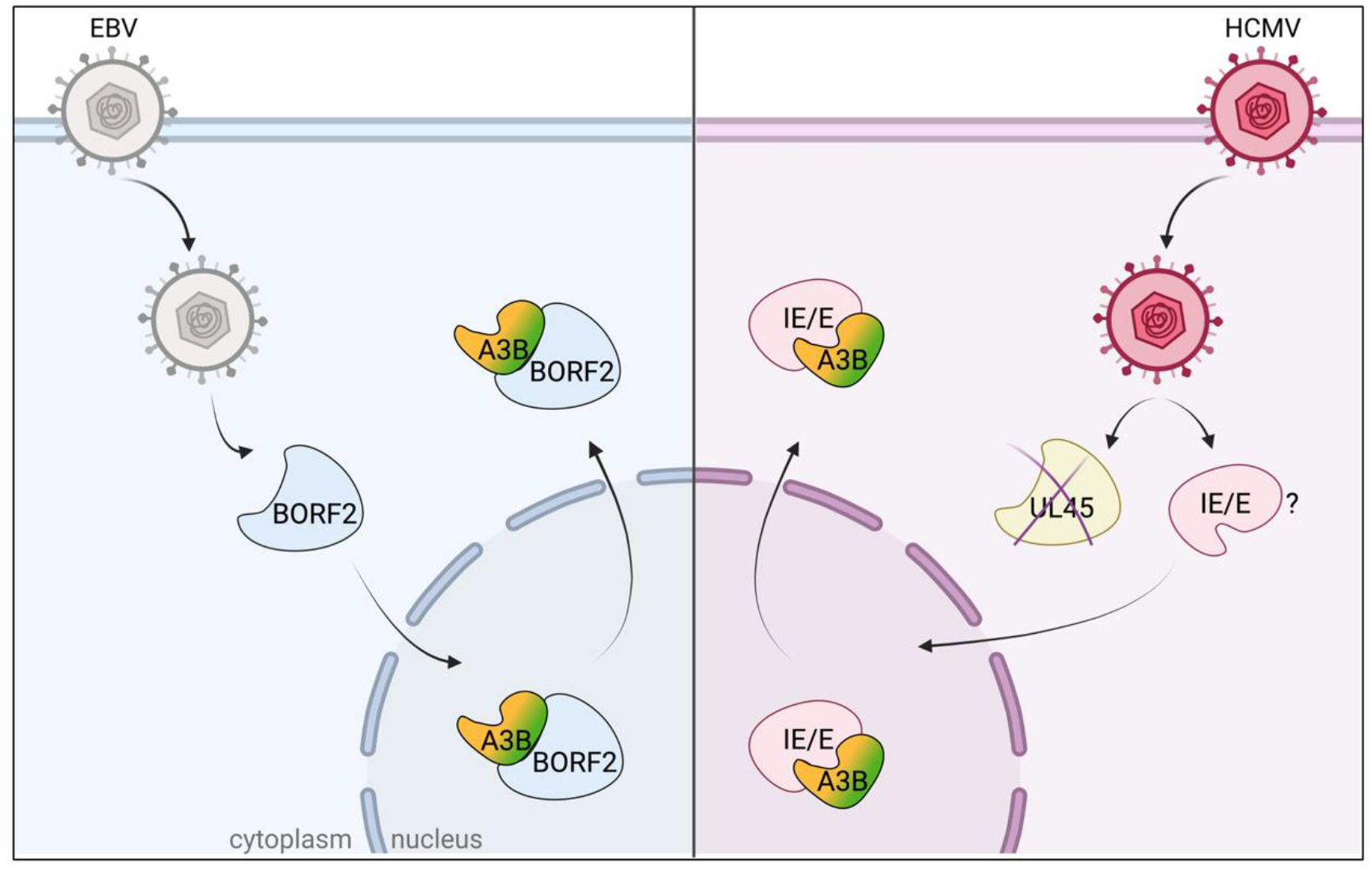
Schematic representation of A3B relocalization mediated by HCMV. Image created with BioRender. EBV (grey), HCMV (magenta), and other herpesviruses (not shown) mediate A3B relocalization from the nucleus to the cytoplasm. EBV utilizes its large RNR subunit (BORF2, light blue) to bind A3B (CTD, green) and promote relocalization. In contrast, HCMV utilizes at least one IE/E protein (pink) to bind A3B (NTD, orange) and promote relocalization. Alternative models are not illustrated for simplicity.

## Discussion

The recent discovery that alpha- and gamma-herpesviruses have evolved strategies to escape from A3-mediated restriction suggested that the beta-herpesvirus HCMV might utilize a similar mechanism to counteract this potent innate immune defense system. Our results demonstrate that HCMV, similar to other herpesviruses, dramatically alters the subcellular localization of the A3B enzyme, relocating it from the nucleus to the cytoplasm. However, this A3B relocalization mechanism is mechanistically distinct, first, by occurring in an RNR-independent manner and, second, by targeting the regulatory N-terminal domain of A3B. In contrast, gamma- and alpha-herpesviruses utilize the large viral RNR subunit to bind to the catalytic C-terminal domain of A3B to mediate relocalization. Moreover, the rapid kinetics of A3B relocalization and pharmacolologic (cycloheximide) and genetic (IE1) requirements described above suggest the involvement of at least one IE/E viral gene product. These results combine to support a working model in which at least one HCMV IE/E protein binds to the regulatory NTD of A3B, promotes its relocalization to the cytoplasm, and thereby protects viral lytic DNA replication intermediates in the nucleus of the cell (**Fig. 7**). Additional studies will be needed to identify the viral factor(s) involved in this process.

A3-mediated restriction of herpesviruses, including HCMV, has been reported (29–32). A3A is upregulated in HCMV-infected decidual tissues and leads to hypermutation of the viral genome (29). Another study reported that A3G is upregulated after HCMV infection of fibroblasts, even if the upregulation does not appear to modulate HCMV replication (31). These studies are certainly interesting, and our work here has not formally excluded these A3s in HCMV restriction. However, given that none of these A3s appear to be counteracted by HCMV (*i.e*., degraded or relocalized), in contrast to A3B described here, they are not likely to pose a significant threat to viral genetic integrity *in vivo*. In contrast, A3B is relocalized away from sites of viral replication by HCMV, which suggests that it may be a *bona fide* threat to the virus during lytic replication. This possibility is supported by the preferred sites of A3B-mediated deamination (5’TC) being depleted from HCMV genomes, consistent with long-term conflicts between this enzyme and HCMV (32, 33). However, this likelihood is difficult to quantify experimentally until the factor(s) involved in A3B neutralization is identified, mutated, and shown to be essential for virus replication in the presence (but not absence) of A3B.

Our studies here add HCMV to the list of herpesviruses that antagonize A3B, suggesting that this function is essential to the success of herpesvirus infection. Our studies are also consistent with the likelihood that this host-pathogen conflict is conserved evolutionarily (8, 11) with ancient origins and remains ongoing to present day. It is surprising, however, that the mechanisms differ on the molecular level in that HCMV (and perhaps other beta-herpesviruses) has evolved a distinct RNR-independent mechanism. If A3B neutralization proves essential for HCMV replication and pathogenesis, it may be possible in the future to drug the neutralization mechanism and enable natural restriction of the infection.

## Materials and Methods

### Cell culture

Cells were cultured at 37°C in a 5% CO2 atmosphere in a Thermo Forma incubator (Thermo Fisher, Waltham, MA). HFF-1 (ATCC, Manassas, VA), U373 (ATCC, Manassas, VA), and HEK293T cells were cultured in DMEM (Cytiva, Marlborough, MA) supplemented with 10% fetal bovine serum (Gibco, Billings, MT), and 1% penicillin/streptomycin (Gibco, Billings, MT). ARPE19 cells (ATCC, Manassas, VA) were cultured in DMEM:F12 media (Gibco, Billings, MT) supplemented with 10% fetal bovine serum (Gibco, Billings, MT) and 1% penicillin/streptomycin (Gibco, Billings, MT). HeLa cells were cultured in RPMI 1640 (Corning) supplemented with 10% fetal bovine serum (Gibco, Billings, MT) and 1% penicillin/streptomycin (Gibco, Billings, MT). All cells were checked periodically for *Mycoplasma* and they always tested negative.

### Viruses and infections

Viruses used in this study were: TB40-mCherry [construction described in (34)]; AD169-GFP [construction described in (35)]; AD169-GFP-ΔUL45 [construction described in (26)]; AD169ΔIE1 [construction described in (36)]. HCMV strain Merlin (GenBank accession NC 006273.2) was purchased from the ATCC (Manassas, VA). The strain FIX and its mutant FIXΔUL45 were a gift by Dr. Elena Percivalle (Fondazione IRCCS Policlinico San Matteo, Pavia, Italy) (37). The TB40-BAC4 and TB40-BAC4-UL45Stop strains used in **Fig. 3E** were produced using a markerless two-step RED-GAM recombination protocol (38, 39). To obtain the BAC of the mutant TB40/E UL45stop the following primers were employed: UL45Stop_Fw: 5’-ATCTACCTGATTTCTTTGTTCTTTTCCTCGTAAACTTATGTAGACTCCGGCTGACGCGGACGAAGGATGACGACGATAAGTAGGG −3’; UL45Stop_Rv: 5’-CCGAGGACACCCGCTGTTCCTCGTCCGCGTCAGCCGGAGTCTACATAAGTTTACGAGGAAAAGCAACCAATTAACCAATTCTGATTAG −3’. All generated recombinant BAC DNAs were controlled for integrity and correctness by sequencing the mutated region. HFF-1 cells were used for the reconstitution of recombinant viruses and virus stock production. Viruses were then propagated by standard procedures as described (40). Briefly, HFF-1 were infected with MOI 0.01 of virus. When robust cytopatic effect (CPE) was observed (between 7 and 14 days) cells were harvested. Then, centrifugation was performed at 15000 g for 30 min. Cell pellets were resuspend in complete media plus 15% Sucrose Phosphate Buffer and sonicated on ice 4X for 10 sec with 15 sec between pulse. Centrifugation was performed at 1300 g for 5 min. Supernatant was collected, aliquoted and frozen at −80°C. The viral titers were calculated using the 50% tissue culture infection dose (TCID50) method upon infection of HFF-1 cells with serially diluted viral supernatants. In all experiments, HFF-1, U373, and ARPE19 were infected with HCMV at an MOI of 3 PFU/cell by diluting the virus into the medium, allowing adsorption for 2 h, and replacing the viral dilution with fresh medium.

### Immunofluorescent microscopy

For immunofluorescence imaging of HCMV infected cells, 5×10^4^ cells/well were seeded in a 24-well plate. After 24 hrs, cells were transduced with lentiviruses encoding for human A3B-HA or A3B-E255A-HA (**Fig. 1A-E**, **Fig. 2A-E**, **Fig. 3D**). 48 hrs after transduction, cells were infected with TB40-mCherry or AD169-GFP for up to 72 hrs as indicated in figure legends. In **Fig. 6B** and **6D**, DMSO, CHX (100 μg/ml), or PAA (100 μg/ml) were added to the virus dilution, and after 2 hrs, when virus was removed, cells were incubated with fresh media and compounds for 24 hrs (CHX) and 48 hrs (PAA). Cells were fixed in 4% formaldehyde for 15 min, permeabilized in 0.2% Triton X-100 in PBS for 10 min, washed three times for 5 min in PBS, and incubated in blocking buffer (2,8 mM KH2PO4, 7,2 mM K2HPO4, 5% goat serum [Gibco, Billings, MT], 5% glycerol, 1% cold water fish gelatin [Sigma, St Louis, MO], 0.04% sodium azide [pH 7.2]) for 1 h. Cells were then incubated with primary rabbit anti-HA (1:2,000) (cat #3724, Cell Signaling, Danvers, MA) or purified rabbit anti-A3B 5210-87-13 [1:300 (25); **Fig. 2G**], or mouse anti-HCMV-IE1 (1:2,000) (cat #MAB810R, EMD Millipore-Sigma, Burlington, MA) (**Supplementary Fig. 2**) overnight at 4°C. Cells were washed 3 times for 5 min with PBS and then incubated with the secondary antibodies goat anti-rabbit IgG Alexa Fluor 488 (1:500) (cat #A11034, Invitrogen, Waltham, MA), or goat anti-rabbit IgG Alexa Fluor 594 (1:500) (cat #A11037, Invitrogen, Waltham, MA), or goat anti-mouse IgG Alexa Fluor 488 (1:500) (cat #A11001, Invitrogen, Waltham, MA) for 2 hrs at room temperature in the dark. Cells were then counterstained with 1 μg/ml Hoechst 33342 for 20 min and rinsed twice for 5 min in PBS.

For immunofluorescence imaging of transfected cells, 5 x 10^4^/well HeLa were transfected with plasmids expressing for 200 ng pcDNA4-BORF2-FLAG, or 200 ng pcDNA4-UL45-FLAG, and 100 ng pcDNA3.1-A3B-HA (**Fig. 3C**). 5 x 10^4^/well ARPE19 were transfected with plasmids expressing for 100 ng pcDNA5TO-A3B-EGFP, pcDNA5TO-A3B-NTD-EGFP, pcDNA5TO-A3B-CTD-EGFP (**Fig. 4A**), and pcDNA4-BORF2-FLAG (**Fig. 4C**). Empty Vector pcDNA3.1 or pcDNA3.1 encoding A3B-HA or other A3x-HA proteins were used in **Supplementary Fig. 1**. After 48 hrs, immunofluorescence was performed as described above. Cells were stained with primary antibodies mouse anti-FLAG (1:2,000) (cat #F1804, Sigma, St Louis, MO) and rabbit anti-HA (1:2,000) (cat #3724, Cell Signaling, Danvers, MA) overnight at 4°C to detect FLAG-tagged RNRs and HA-tagged A3B, respectively. Goat anti-mouse IgG Alexa Fluor 488 (1:500) (cat #A11001, Invitrogen, Waltham, MA) and goat anti-rabbit IgG Alexa Fluor 594 (1:500) (cat #A11037, Invitrogen, Waltham, MA) were used as secondary antibodies.

Images were collected at 20x magnification using an EVOS FL Cell Imaging System (ThermoFisher Scientific). Quantification was performed using Image J software, counting the percentage of cells with relocalized A3B or the ratio of nuclear/cytoplasmic A3B. Quantification was performed counting cells from n=3 independent experimental replicates. GraphPad Prism 9 was used to prepare graphs and statistical analyses (unpaired student’s t test).

### Coimmunoprecipitation experiments

HEK293T (2.5 x 10^5^/well) cells were grown in 6-well plates and transfected with pcDNA3.1 plasmids encoding human A3A-HA, A3B-HA, and A3G-HA together or not with pcDNA4-BORF2-FLAG or pcDNA4-UL45-FLAG, and 6 μl TransIT-LT1 (Mirus, Madison, WI) in 200 μl serum-free Opti-MEM (Thermo Fisher Scientific, Waltham, MA). After 48 h, whole cells were harvested in 300 μl of ice-cold lysis buffer (150 mM NaCl, 50 mM Tris-HCl, 10% glycerol, 1% IGEPAL [Sigma, St Louis, MO], and cOmplete EDTA-free protease inhibitor cocktail [Roche] [pH 7.4]). Cells were vortexed, incubated on ice for 30 min, and then sonicated. Whole-cell lysate (30 μl) were aliquoted for input detection. Lysed cells were centrifuged at 13,000 rpm for 15 min to pellet debris, and the supernatant was resuspended with 25 μl anti-FLAG M2 magnetic beads (Sigma, St Louis, MO) for overnight incubation at 4°C with gentle rotation. Beads were washed three times in 700 μl of lysis buffer. Bound protein was eluted in 30 μl of elution buffer (0.15 mg/ml 3xFLAG peptide [Sigma, St Louis, MO] in 150 mM NaCl, 50 mM Tris-HCl, 10% glycerol, and 0.05% Tergitol [pH 7.4]). Input and eluted proteins were analyzed by western blot. Membranes were stained with mouse anti-FLAG (1:5,000) (cat #3724, Sigma, St Louis, MO), mouse anti-tubulin (1:10,000) (cat # T5168, Sigma, St Louis, MO), and rabbit anti-HA (1:3,000) (cat #3724, Cell Signaling, Danvers, MA). After washing, membranes were incubated with an anti-rabbit IgG horseradish peroxidase-conjugated (HPR) secondary antibody (1:10,000) (cat #211032171, Jackson ImmunoResearch, West Grove, PA) and an anti-mouse IRDye 800CW (1:10,000) (cat #C70919-05, LI-COR, Lincoln, NE) (**Fig 3A**).

### DNA deaminase activity assays

HEK293T (5 x 10^5^/well) cells were seeded into 6-well plates and, after 24 hrs, transfected with 200 ng pcDNA4-BORF2-FLAG, or 200 ng pcDNA4-UL45-FLAG, and 100 ng pcDNA3.1-A3B-HA. After 48 hrs, cells were harvested, resuspended in 100 μl of reaction buffer (25 mM HEPES, 15 mM EDTA, 10% glycerol, 1 tablet of Sigma-Aldrich cOmplete Protease Inhibitor Cocktail), and sonicated at the lowest setting. Whole-cell lysates were then centrifuged at 10,000 x g for 20 min. The clarified supernatant was incubated with 4 pmol of oligonucleotide (5’-ATTATTATTATTCAAATGGATTTATTTATTTATTTATTTATTT-fluorescein), 0.025 U uracil DNA glycosylase (UDG), 1x UDG buffer (NEB), and 1.75 U RNase A at 37°C for 2 hrs. Deamination mixtures were treated with 100 mM NaOH at 95°C for 10 min. Samples were then separated on 15% Tris-borate-EDTA-urea gel. Fluorescence was measured using a Typhoon FLA-7000 image reader (**Fig. 3B**).

## Acknowledgements

We thank members of the Harris laboratory for support and constructive feedback. This work was supported by NIAID R37-AI064046 and NCI P01-CA234228. Salary support for AZC was provided in part by NIH training grants F30-CA200432 and T32-GM008244. Salary support for AAA was provided in part by NIH T32-AI83196 from the University of Minnesota’s Institute for Molecular Virology Training program. Salary support for SNM was provided by NIAID F31-AI161910 and subsequently an HHMI Gilliam Fellowship. RSH is an Investigator of the Howard Hughes Medical Institute, a CPRIT Scholar, and the Ewing Halsell President’s Council Distinguished Chair. The authors have no competing interests to declare.

## Contributions

EF and RSH conceptualized the study. EF, AZC, AAA, and BS performed experiments. EF curated the data, generated figures, and was responsible for formal data analyses. EF and RSH wrote the initial draft of the paper and all authors contributed to revisions. SNM, JRL, CJB, WAB, VDO, MB, and RSH provided resources. RSH was responsible for funding acquisition.

